# Distribution and dynamics of chromatin states in the *C. elegans* germ line

**DOI:** 10.1101/2023.05.11.540452

**Authors:** Mariateresa Mazzetto, Lauren E. Gonzalez, Nancy Sanchez, Valerie Reinke

## Abstract

Chromatin organization in the *C. elegans* germ line is tightly regulated and critical for germ cell differentiation. While certain germline epigenetic regulatory mechanisms have been identified, how they influence chromatin structure and ultimately gene expression remain unclear, in part because most genomic studies have focused on data collected from whole worms. We therefore analyzed publicly available histone modification and chromatin accessibility data from isolated undifferentiated germ nuclei to define chromatin states. We then correlated these states with overall transcript abundance, spatio-temporal expression patterns, and the function of small RNA pathways. Because the essential role of the germ line is to transmit genetic information to the next generation and establish gene expression in the early embryo, we compared epigenetic and transcriptomic profiles from undifferentiated germ cells, oocytes, and embryos to define the epigenetic changes during this developmental transition. The active histone modification H3K4me3 exhibits particularly dynamic remodeling as germ cells differentiate into oocytes. Our results highlight the dynamism of the chromatin landscape in germ cells, and provide a resource for future investigation into epigenetic regulatory mechanisms.

## INTRODUCTION

Epigenetic mechanisms are essential to correctly establish tissue-specific gene expression programs during development. At the genomic level, such mechanisms include deposition of histone modifications and histone variants that lead to alterations in chromatin structure and organization, which can have both global and local effects on gene expression. Small RNA pathways are also mediators of epigenetic information by influencing transcript stability as well as feeding back to the nucleus to affect chromatin states at target loci.

Epigenetic mechanisms are especially critical in germ cells, which have the task of transmitting both genetic and epigenetic information from parent to progeny. In *C. elegans*, multiple epigenetic mechanisms are implicated in different aspects of germ cell differentiation and function. These mechanisms include the mutually exclusive relationship of the repressive H3K27me3 and active H3K36me3 histone modifications, which reinforce permissive chromatin environments on autosomes [19,34,38], and enrich repressive marks on the X chromosome, leading to its inactivation [6,54]. The relationship of these two histone modifications with other modifications in the germ line is poorly understood. Additionally, several distinct small RNA pathways are very active in germ cells, and affect the stability of target transcripts as well as select histone modifications at target loci [18,48]. However, the extent to which target gene chromatin state is altered as a consequence of the activity of individual small RNA pathways has not been systematically determined. Finally, while epigenetic information is thought to be critical for highly differentiated gametes to reactivate totipotency in the zygote and initiate successful embryogenesis in the absence of active transcription, the accompanying changes to chromatin structure and how they contribute to correct genome activation in the early embryo, are not well characterized [44].

Functional and genomic studies have revealed some epigenetic regulatory mechanisms that influence germline chromatin organization and establish reproducible gene expression programs. However, the field has been limited by the fact that most genomic studies in *C. elegans* have been performed on whole animals, which simultaneously captures both germline and somatic tissues and therefore obscures germline-specific mechanisms. We recently addressed this limitation by developing a protocol to collect isolated germ nuclei (IGN) at sufficient numbers that allow for genomic assays [23]. Over the last few years, we and others have successfully used this protocol for different high-throughput sequencing assays in IGN, such as transcriptome profiling (RNA-seq), histone modification profiling via Chromatin Immunoprecipitation (ChIP-seq), and chromatin accessibility profiling via Assay for Transposon-Accessible Chromatin (ATAC-seq) [45].

In this study, we characterize *C. elegans* chromatin regulatory mechanisms in IGN using these datasets. We first define diverse chromatin states in the germline genome, reveal intra- and inter-chromosomal patterns of histone modifications and chromatin accessibility, and identify chromatin states that are unique to the germ line relative to the soma. To observe how chromatin states at individual genes correlate with spatial and temporal gene regulation during germline differentiation, we examined genes with preferential expression in pregametic cells, spermatogenesis, or oogenesis [31], as well as genes targeted by multiple small RNA pathways [48]. Together, these analyses revealed unexpected disconnects between active histone modifications, open chromatin, and transcript abundance, as well as distinct effects by different small RNA pathways on chromatin state of target genes. We also examined how chromatin status changes during gametogenesis and the maternal-to-zygotic transition, and found that one histone modification, H3K4me3, exhibits particularly dynamic remodeling as germ cells differentiate into oocytes. Overall, these analyses improve our understanding of the dynamic chromatin state found in germ cells, and provide a platform for continued investigation into global and local epigenetic germline regulatory mechanisms.

## METHODS

### Datasets and genome versions

Throughout this study, we used the WBcel235/ce11 (WS235) version of the *C. elegans* genome, and WormBase WS286 genome annotations - with coordinates backlifted to WBcel235/ce11 (WS235).

Datasets used in these analyses were all extracted from published work:

- H3K4me3 and H3K27ac ChIP-seq datasets from IGN from wild type (strain VC2010) and *glp-1(allele?)* mutants at the young adult stage used in this analysis refer to accession number GSE117061 [23], whereas H3K36me3 and H3K27me3 ChIP-seq datasets refer to GSE147401 [37];
- H3K9me3 data from IGN samples refer to GSE232141 [45], whereas data from *glp-1* young adult mutants refer to GSM3141347 [46];
- ChIP-seq datasets from mixed embryos refer to GSE114440 [25], GSM3148388 [8], and GSE168923 [13];
- ChIP-seq datasets from early embryos, sperm and oocytes were all obtained from GSE115709 [51];
- Pol II whole worm ChIP-seq data were acquired from GSE162063 [15];
- ATAC-seq data from IGN were taken from GSE232139 [45], whereas ATAC-seq data from mixed embryos and *glp-1* mutants at young adult stage were taken from GSE114440 [25];
- RNA-seq data from IGN and *glp-1* mutants acquired at the young adult stage were obtained from GSE117061 [23].

### ChIP-seq and ATAC-seq data processing

Since the data used in this study came from many different sources, we re-processed all data identically as follows. The raw data in FASTQ format were aligned to the reference genome ce11 using Bowtie2 [29], and then converted to SAM, sorted, and filtered for uniquely aligned reads (-q 10) using SAMtools [33]. Peaks were called with MACS2 [15] using the standard options (-q 0.001 –nomodel – extsize 150). BigWig files were obtained using the bigwigCompare function in the Deeptools software [42] and read-normalized on input files.

Metagene plots were obtained using the computeMatrix and plotProfile tools in Deeptools using the reference-point mode (--regionBodyLength 3000 --beforeRegionStartLength 2000 -- afterRegionStartLength 2000 --binSize 50).

Read counting was performed using the dbacount function in the DiffBind package [55]; Annotation of genomic regions and gene ontology enrichment were performed using the ChIPSeeker and ClusterProfiler packages [56, 57].

To visualize the distribution of histone marks along the chromosomes we used the rtracklayer package for importing all bigwig files [30].

### Annotation of chromatin states

Annotation of chromatin states was performed on histone marks (ChIP-seq) and chromatin accessibility (ATAC-seq) data using the ChromHMM software [4]. The program partitions the genome in 200-bp intervals, then assesses the presence or absence of a specific mark on each bin, and uses a hidden Markov model on the resulting calls to learn a chromatin-state model, and finally, obtains the annotation of each state occurrences on the genome. The standard output contains a mark emission table containing the probability of each mark in a specific state.

States were named on the basis of the combination of marks visualized, the probability of each state at annotated genomic regions, and the distribution relative to the transcription start and end sites (TSS and TES, respectively) [16].

The analysis was done on both germ line and somatic data using the “independent” mode. We tested multiple numbers of states and selected a 12-state model for all further analyses, which best represented the diverse relationships between patterns of histone marks, open chromatin, and genomic features.

### Data visualization

To visualize our results the following R and Python packages were used:

- Gviz and KaryotypeR packages for the chromosomal views of chromatin states [20,24];
- The Venneuler package for Venn Diagrams;
- The Deeptools package for epigenetic profile plots around specific genes (both metagene plots and heatmaps), with the computeMatrix, plotProfile and plotHeatmap functions [42];
- The ComplexHeatmap package for heatmaps showing the average signal of epigenetic marks around each chromatin state, as well as embryo-IGN correlations [22];
- The ReactomePA package for plotting GO results [58].

### Gene lists

Genes with enriched expression in sperm, oocytes, and pregametic cells (Figure 2) were taken from Lee et al. [31], whereas sperm genes targeted by the piRNA pathway were obtained from Cornes et al [11]. Genes with tissue-specific expression (Figure 4) were acquired from Serizay et al. [47]. Finally, the lists of small RNA pathway target genes were taken from Seroussi et al. [48].

## RESULTS

### Characterization of germline chromatin states

To characterize the distribution of regulatory states of *C. elegans* germline chromatin in wild type adult hermaphrodites, we analyzed epigenetic data from isolated germ nuclei (IGN), which primarily comprise undifferentiated proliferating and early meiotic germ cells [23]. Datasets include profiles of five well-characterized histone marks (H3K27me3, H3K9me3, H3K36me3, H3K4me3, and H3K27ac), transcript abundance, and chromatin accessibility data (Methods). To identify active or inactive genome regions that correlate with particular combinations of histone modifications and associate these data with genomic features such as promoters, enhancers, exons and introns, we annotated chromatin states across the genome using ChromHMM [16]. This program discovers chromatin state signatures using a hidden Markov Model (HMM) that measures the presence or absence of each mark, singly or in combination, across genomic features to define functionally distinct chromatin states. ChromHMM also provides the quantified distribution (“coverage”) of each state across the genome. We defined 12 chromatin states characterized by specific combinations of histone marks and distinct distributions relative to genomic features (Figure 1A, Figure S1A-B). Six of the states represent repressed or inactive chromatin. We named two states with high levels of H3K9me3 alone or H3K9me3 together with H3K27me3 as “heterochromatin”; a state with high levels of H3K27me3 alone as “PC (Polycomb) repressed”; a state with low levels of any mark as “quiescent”; and two states with high chromatin accessibility but the presence of H3K27me3 signal around the TSS as “inactive TSS” (Figure 1A). For the six states representing active chromatin, we defined one state with accessible chromatin and both active and repressive marks distributed around the TSS as “primed TSS”; two states with combinations of active marks near the TSS as “active TSS”; one state with high levels of H3K36me3 and H3K4me3 on gene bodies as “active transcription”; and two states with combinations of active marks, distribution on the gene body, and accessible chromatin as “euchromatin”. As expected, genes associated with active chromatin states (states 7-12) were characterized by higher transcript abundance (Figure 1B) and transcript integrity (Figure 1C), while transcripts from genes in inactive or repressive chromatin states were less abundant and more prone to degradation. In addition, Gene Ontology (GO) analysis for the genes contained in each chromatin state were enriched for developmental and germ line-related categories in active states, neuronal-related categories in inactive states, and, uniquely in heterochromatin, glycoprotein metabolism (Figure S1C). These observations show that germline chromatin can be segmented into regulatory states with distinct features that reflect known gene function and gene expression in that tissue.

**Figure 1.**
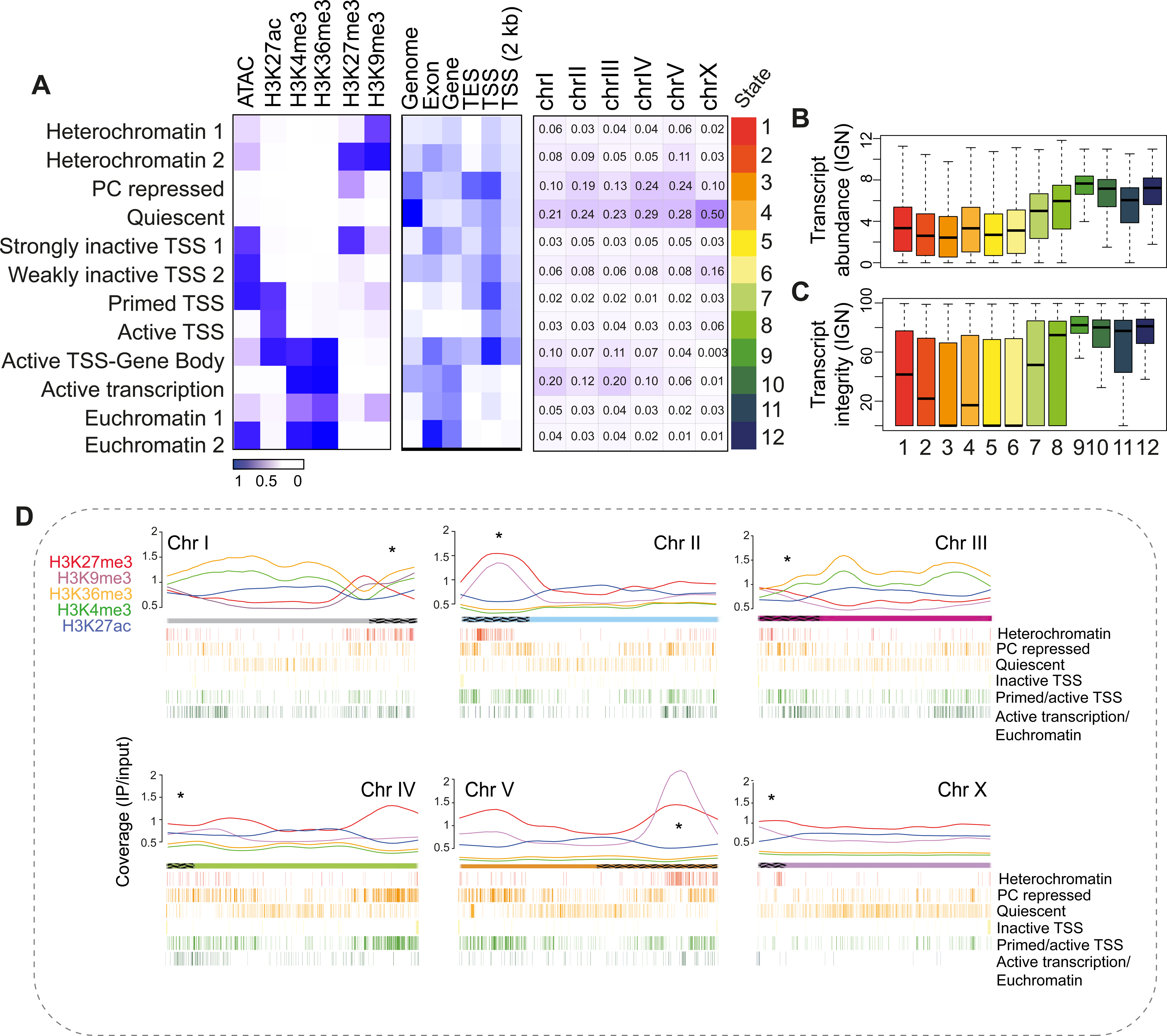
Characteristics of germline chromatin state. A) Annotated chromatin states from isolated germ nuclei (IGN). Left panel: the probability of each histone mark or open chromatin for each of the 12 defined states (as a range from 0 to 1), are plotted as a heatmap. Middle panel: the genomic coverage across gene features for each state. Right panel: percentage of each chromosome in each state. B) Transcript abundance (as log FPKM) of the genes within each chromatin state. C) Percent transcript integrity of the genes within each chromatin state. D) Distribution of individual histone modifications along each chromosome, plotted as the normalized signal (IP/input ratio). Chromatin states with similar distribution of histone modifications were combined into groups and displayed.

In the germ line, autosomes are generally well-expressed and enriched with active marks, while the X chromosome is generally silenced, with enriched H3K27me3 and low levels of active marks [6,19,34,38,54]. Additionally, meiotic recombination events are concentrated toward the centers of autosomes, but are broadly distributed along the length of the X chromosome [3,35]. We compared chromatin state and histone modification patterns to identify both inter- and intra-chromosomal differences at higher resolution. The coverage of each chromatin state relative to percent chromosome length (Figure 1A, right panel) demonstrates that the autosomes display functionally diverse (inactive, quiescent, and high transcription) states. By contrast, the X chromosome displays a quiescent or inactive state for 84% of its length, a scattered distribution of heterochromatin states, low levels of active marks, and higher levels of the H3K27me3 mark compared to active marks (Figure 1D).

We also observed distinct intra-chromosomal patterns of chromatin states and histone modification distributions (Figure 1D). While heterochromatin is preferentially concentrated on chromosomal arms, euchromatin is mainly distributed on the central part of each chromosome, which are generally gene-dense and enriched for essential genes [9,26]. In addition, heterochromatin states and repressive histone marks, especially H3K27me3, tend to be distributed around the meiotic pairing center, while active marks are depleted, as previously reported [3,9,40]. In addition to these general, expected patterns, each chromosome has unique aspects of chromatin state distribution. For example, chromosome III has relatively low levels of repressive marks across the whole chromosome length, even on the chromosomal arms. Additionally, chromosome IV does not show specific changes in levels of histone marks at the pairing center, perhaps due to having relatively few pairing center sequence motifs [35,39]. Chromosome IV also displays high levels of H3K27me3 at the right arm of the chromosome, which is the site of a 3 Mb domain containing thousands of tiny genes encoding type I piRNAs. Enrichment of H3K27me3 on genes encoding this class of small RNAs was previously observed in whole animal ChIP-seq analysis [4]. Finally, a notably strong H3K9me3 signal on Chromosome V correlates with a genomic region enriched for pseudogenes and G protein-coupled receptor genes with neuron-restricted expression.

With these observations, we show that annotation of chromatin states using purified germ nuclei gives a comprehensive characterization of chromatin regulatory states among and within chromosomes in the *C. elegans* germline.

### Characterization of somatic chromatin states relative to the germ line

To define unique aspects of the germline chromatin landscape, we wished to contrast chromatin states with those in somatic tissues. We therefore computed chromatin states for adult worms lacking a germline and composed only of somatic tissues, using published epigenetic and transcriptomic data representing the same chromatin features analyzed in the germ line [23,25]. We again classified 12 chromatin states characterized by specific combinations of histone marks and distinct distributions along the genome (Figure 2A, S2A). Similar to the germline, genes associated with active chromatin states in the soma were characterized by higher transcript abundance (Figure 2B) and transcript integrity (Figure 2C), while genes contained in quiescent or repressive chromatin states were less abundant and more prone to degradation. Unlike the germ line, the X chromosome in the soma is not enriched for inactive states (Figure 2A, S2A). Moreover, the levels of active marks are generally lower along all chromosomes in the soma, while H3K27me3 is particularly high not only on the meiotic pairing center but over the length of each chromosome, with the exceptions of chromosomes I and III (Figure S2B). High levels of H3K27me3 might be the result of mixing many different cell types, in which most tissue-specific genes would be turned off in most tissues, leading to a higher H3K27me3 signal on average along the chromosome. Gene Ontology analysis revealed an enrichment in active stages for developmental and proliferation categories, gene categories that likely span multiple tissue types; heterochromatin regions instead showed an enrichment for chemosensory pathways and immune response categories, which are likely only involved in a few specific cell types and therefore need to be repressed in most cell types of the worm (Figure S2C). Overall, these chromatin states from whole somatic tissue provide a useful platform to contrast with germline chromatin states.

**Figure 2.**
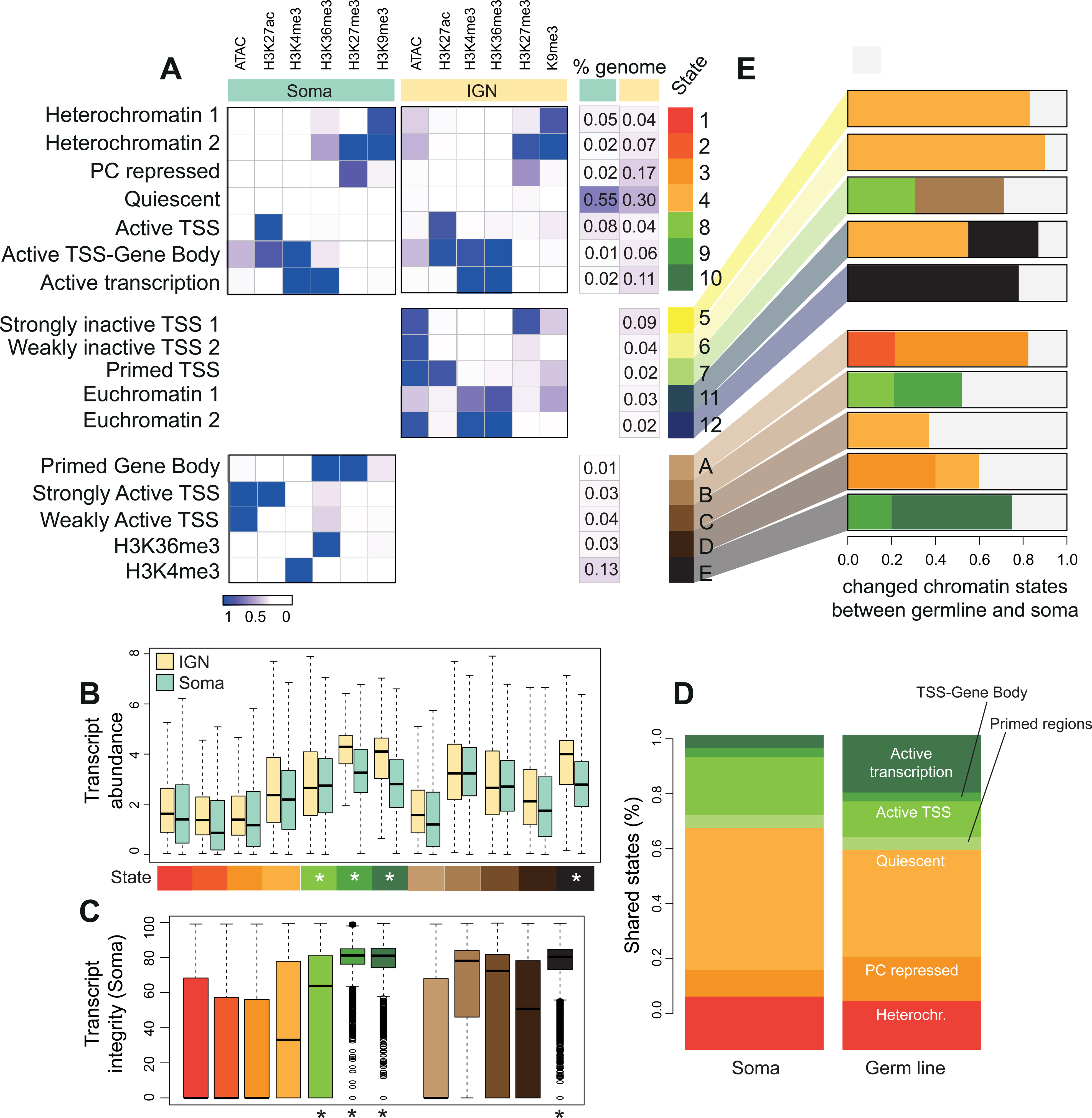
Chromatin states unique to germ line and soma. A) Annotated chromatin states from adult mixed somatic cells compared to germline states from Figure 1, divided into “shared” states (above), “only-IGN” states (center), and “only-soma” states (below). The probability of each histone mark or open chromatin for each of the 12 defined states (as a range from 0 to 1), are plotted as a heatmap. with percentage of each chromosome in each state at left. B) Transcript abundance (as log FPKM) of the genes located in each chromatin state. Asterisks mark active states. C) Percent transcript integrity of the genes located in each chromatin state. D) Percent sequences in selected chromatin states in both IGN and soma. E) Examination of chromatin state switching for soma and germline-specific states. For genomic regions found in a tissue-specific chromatin state, the opposite tissue was examined for changes to chromatin state. Only chromatin states with a genomic coverage > 20% were plotted.

We therefore directly compared chromatin states between the germ line and soma to identify germline-specific aspects of chromatin regulation. Seven somatic chromatin states are very similar to those in the germ line: two heterochromatin states, as well as the PC repressed, quiescent, active TSS, active TSS-Gene Body, and active transcription states, which together represent 75-80% of the genome in both the soma and germ line. Overall, about 58% of the genome retained the same state between germline and soma, with specific chromatin states differing in percent genome coverage (Figure 2D). Additionally, both the germ line and soma exhibit unique chromatin states. For example, in the germ line about 9% of the genome exhibits the silencing mark H3K27me3 at promoters in sites of open chromatin (strongly inactive TSS state 5), a state that is not found in the soma. By contrast, 16% of the genome in the soma has a single active histone mark (H3K36me3 or H3K4me3, states D and E) while those two marks are always found together in the germline chromatin states.

We then investigated how chromatin states changed between germ line and soma at specific regions of the genome (Figure 2E). As an example, regions marked by the germline-specific “strongly inactive TSS” state 5 are found largely in the quiescent state in the soma. By contrast, regions marked by the “H3K36me3 only” state D in the soma lose H3K36me3 and become quiescent (state 4) or PC repressed (state 3) in the germline, while regions in the “H3K4me3 only” state E gain H3K36me3 and become the active states 9 (TSS-Gene Body) and 10 (active transcription) in the germline. At the resolution of individual genes, we also found many changes between germline and somatic epigenetic profiles (Figure S2D-F). For instance, *bath-4*, a germline-specific gene enriched in germline precursor cells, shows strong enrichment for active marks in the germline but enrichment for H3K27me3 in the soma. Notably, H3K36me3 and H3K27me3 are mutually exclusive in the germ line, but show some overlap at individual genes in the soma (Figure S2G), which could again be related to mixing multiple cell types. Indeed, germline-specific genes were poorly expressed in the soma, whereas tissue-specific somatic genes were more highly transcribed (Figure S2H). In sum, comparing germ line chromatin states with those found in the soma reveals both shared and distinct epigenetic profiles, suggesting that there are unique aspects to epigenetic regulation and/or function in the germ line.

### Using chromatin states to understand germ line regulatory mechanisms

The *C. elegans* germ line is characterized by an organized trajectory along which proliferating germ cells enter meiosis I and either differentiate into sperm at the L4 stage or into oocytes as adults. These temporal and spatial dynamics result in extensive differential gene regulation [28,43,49] that defines groups of genes with enriched expression prior to differentiation (“pregametic”), during spermatogenesis or during oogenesis [31]. To better understand the relationship between chromatin states, gene regulation, and germline differentiation, we examined the chromatin state distribution specifically associated with these gene sets.

Of the three groups of genes, the pregametic genes are best represented by IGN, which primarily captures nuclei from morphologically undifferentiated germ cells in the distal proliferative and medial pachytene regions of the adult germline [23]. We found that 70% of pregametic genes are in states 9 and 10, “active TSS-gene body” and “active transcription”, with high levels of H3K36me3, H3K4me3, and H3K27ac (Figure 3A, S3A) and high transcript abundance (Figure 3C). Surprisingly, these genes do not have the most open chromatin; instead states 7 and 8, “primed TSS”, and “active TSS”, which represent only 4% of the pregametic genes, have substantially more open chromatin (Figure 3B). This result suggests that many genes can be abundantly expressed even if the chromatin is not maximally accessible. Strikingly, 16% of pregametic genes are associated with inactive states 1-6, yet show at least some level of expression, with particularly high transcript abundance for state 1, which is characterized by both H3K9me3 and H3K27me3 (Figure 3C). We hypothesized that the high transcript abundance in these inactive states could be due to post-transcriptional stabilization of mRNAs produced during larval germline development, rather than ongoing transcription. To test this hypothesis, we looked at whole worm RNA Polymerase II signal [15] around pregametic genes in each chromatin state. Genes in active states show high levels of Pol II occupancy across the gene body, whereas genes in inactive states, including state 1, display low Pol II signal (Figure S3B). Together, these analyses suggest that pregametic genes with repressive chromatin marks might no longer be actively transcribed in the adult germ line, but transcript levels remain high because of post-transcriptional stabilization of pre-existing mRNAs.

**Figure 3.**
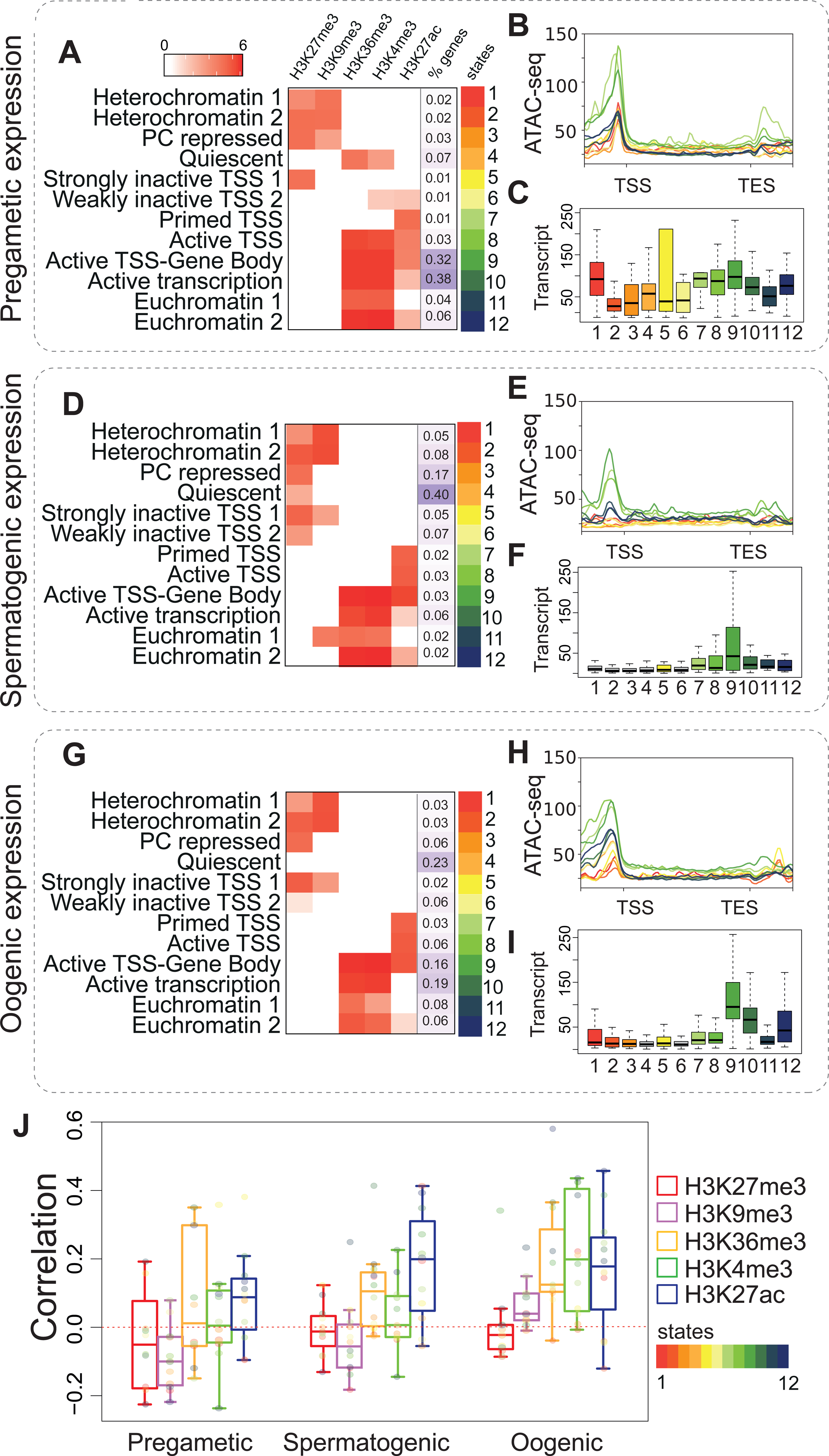
A-I) Chromatin states of classes of germline-expressed genes in IGN. (A-C) pregamete-enriched gene class, (D-F) spermatogenesis-enriched gene class, and (G-I) oogenesis-enriched gene class [31]. (A,D,G) Heatmap of histone mark coverage in each state, with percent chromosomes in each state. (B,E,H) Chromatin accessibility across chromatin states. Each line represents a chromatin state. Normalized signals (as IP/input ratio) were plotted. Upstream region (TSS-2000bp), gene body, and downstream region (TES+2000 bp) are included. (C,F,I) Transcript abundance as FPKM. Values related to each state are plotted. J) Pearson correlation between transcript abundance (FPKM) in IGN and histone mark levels (as normalized IP/input signal) for spermatogenic, oogenic, and pregametic gene classes. Each point represents a single chromatin state with color code as in Figure 1.

We then focused on how the spermatogenesis gene class is regulated in adult IGN. These genes are expected to be inactive since spermatogenesis stops abruptly at the end of L4 upon the switch to oogenesis in hermaphrodites [43]. As expected, 82% of spermatogenic genes are found in inactive chromatin states 1-6, all of which have moderate to high levels of H3K27me3, suggesting that this mark is a key component involved in silencing these genes (Figure 3D, Figure S4A). In addition, genes involved in spermatogenesis are mostly in a closed chromatin configuration (Figure 3E) with low transcript abundance (Figure 3F). Recently, the piRNA pathway [52] was demonstrated to downregulate spermatogenesis-expressed genes in the adult germ line, dependent on the nuclear Argonaute protein HRDE-1 [11]. HRDE-1 is expected to recruit H3K9me3 to target genes, although this outcome was not directly tested. Only 13% of all spermatogenesis genes are associated with H3K9me3 (states 1 and 2), but 30% of the subset that are piRNA target genes are in states 1 and 2 (Figure S3B). The piRNA pathway also targets pseudogenes, and we found that 12 of 23 pseudogenes with spermatogenesis-enriched expression are in states 1 and 2, a four-fold enrichment. Together, these results support the idea that at least a subset of piRNA target genes expressed during spermatogenesis are likely to be silenced through recruitment of H3K9me3.

Unexpectedly, spermatogenesis genes in state 9 “active TSS-Gene Body” show high transcript abundance, suggesting that they were still expressed or stabilized in adult germ cells. We found that genes in this and similar states do not show a spike in expression at the L4 stage, as seen for other spermatogenesis genes, but instead have consistent expression across development (Figure S4C) as well as similar levels between hermaphrodite soma and males at the L4 stage (Figure S4D), indicating that their expression is not restricted to spermatogenesis but occurs in multiple tissues. Consistent with this possibility, Gene Ontology analysis revealed that these genes are enriched for functions in basic metabolism (Figure S4E), and thus are likely to be broadly expressed.

Genes in the oogenesis gene class are distributed more evenly among chromatin states than either the pregametic or spermatogenesis groups, with 43% in inactive states (1-6) and 57% in active states (7-12) in adult IGN (Figure 2G and Figure S5A). We speculated that oogenesis genes in inactive chromatin states might not become expressed until oocytes start to form, while genes in active states are expressed in IGN. To explore this possibility, we examined oogenesis genes on the X chromosome, which are well-characterized as being specifically upregulated in developing oocytes in the proximal germline [26,52]. A total of 60% of X-linked oogenesis genes are in the inactive states 1-6, with 34% found in state 4 “quiescent”, which has very low levels of any modification (Figure S5B). Indeed, almost half (47%) of oogenesis genes in the quiescent state are located on the X. However, 40% of X-linked oogenesis genes are in active states 7-12. Thus, even though the expression of most X-linked oogenesis-enriched genes is restricted to the proximal germline, they can exhibit active, quiescent, or inactive states in IGN. Specific examples highlight the heterogeneity found in oogenesis genes (Figure S5C-E). For genes found in states 8 (“Active TSS”), 9 (“Active TSS-Gene Body”), 10 (“High transcription), and 12 (“Euchromatin”), Gene Ontology analysis revealed an enrichment of for categories related to gamete generation and reproduction (Figure S5F); however, overall a wide variety of gene functions are found in the different chromatin states. Together, these observations point to diverse regulatory mechanisms beyond chromatin state that combine to determine expression levels of genes in the oogenesis-enriched class.

We also noted common trends for all three groups of genes with germline-enriched expression. Chromatin state 7 “primed TSS”, which represents only 1-3% of genes in each group and is specifically marked by H3K27ac only, consistently has the most open chromatin conformation (Figure 3B, E, H). However, open chromatin does not directly correlate with transcript abundance, as genes in this state typically have lower transcript abundance compared to other active states with less open chromatin (Figure 3C, F, I). Instead, peak transcript abundance consistently occurs in state 9 “active TSS-gene body”, which is characterized by high levels of all three activating marks H3K36me3, H3K4me3, and H3K27ac with only an intermediate level of open chromatin. Overall, transcript abundance in all three categories is most correlated with H3K27ac, while the expression of oogenesis-expressed genes in IGN is additionally positively correlated with H3K36me3 and H3K4me3 levels (Figure 3J). In sum, diverse chromatin environments in IGN at genes with spatial and temporal regulation during germ cell differentiation reveal complex relationships between chromatin state and transcript abundance and support the role of extensive post-transcriptional regulation.

### Chromatin state of gene targets of small RNA pathways

The disconnect between chromatin state and transcript abundance for many of the pregametic, spermatogenesis, and oogenesis-expressed genes suggests extensive post-transcriptional regulation. In *C. elegans*, diverse small RNA pathways are key post-transcriptional mediators of gene regulation, especially in the germ line. These germline small RNA pathways typically affect transcript stability, but some have also been implicated in altering chromatin state at target genes as a downstream effect [18]. However, whether particular states are predominant for specific pathway target genes has not been systematically investigated specifically in germ cells. Classes of small RNAs are primarily defined by their associated Argonaute (AGO) protein, which dictates their subcellular localization and accessibility to targets [14]. A recent study defined four distinct groups of AGO proteins based on shared targets and overlapping functions and phenotypes: the germline-specific CSR-1 and WAGO-1 groups, the sperm-enriched ALG-3/4 group, and the somatic ERGO-1 group [48].

To investigate the relationship between epigenetic state and these functionally distinct AGO/small RNA pathways, we first examined the distribution of germline chromatin states around the gene targets of each AGO in each subgroup. We found that ∼80% of the targets of each AGO in the CSR-1 group (CSR-1, VRSA-1, and WAGO-4) are associated with active chromatin states, as expected given that the CSR-1 pathway primarily targets germline-expressed genes to promote their transcript levels (Figure 4A, right). However, AGOs in the ALG-3/4 group (ALG-3, ALG-4, RDE-1, and WAGO-10) have a more variable distribution, with ALG-3/4 target genes distributed roughly evenly between active and inactive chromatin states, while almost 80% of RDE-1 and WAGO-10 target genes are associated with inactive chromatin states (Figure 4A, left). Targets of the ERGO-1 group, which primarily represent somatic genes not expressed in the germline [47], are mostly associated with inactive states (Figure S6A). Similarly, AGOs in the WAGO-1 group are important for the repression of transposable elements [48], and the gene targets are mostly in inactive chromatin states (Figure S6A). Target genes of the CSR-1 and ALG-3/4 groups have higher transcript abundance in IGN compared to the soma, while the targets of the ERGO-1 and WAGO-1 groups show similar levels of transcript abundance in IGN and soma (Figure 4B, Figure S6B).

**Figure 4.**
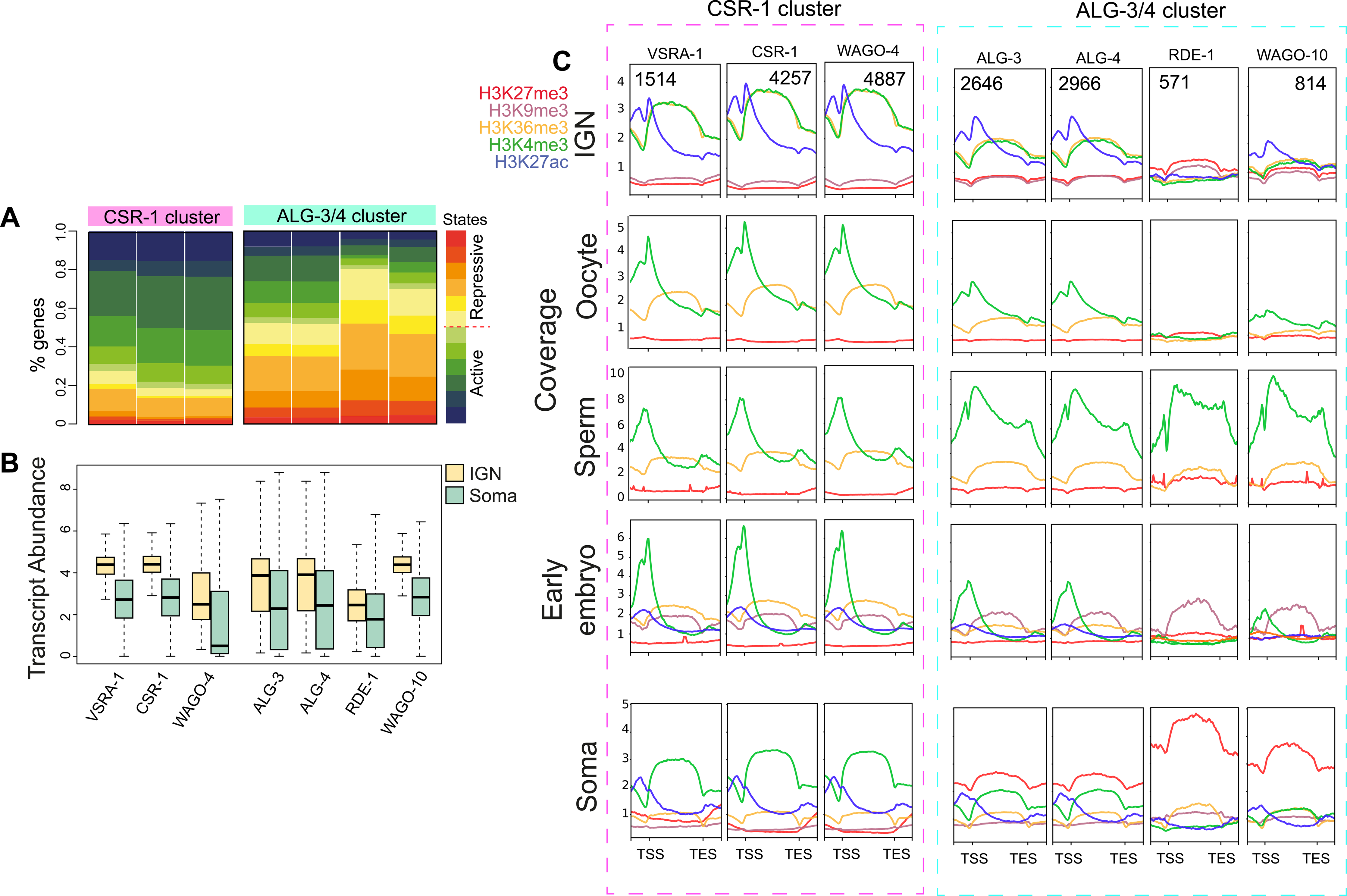
Chromatin states of AGO target genes. A) Distribution of chromatin states along targets of AGO proteins in the CSR-1 and ALG-3/4 groups [48]. B) Transcript abundance of these targets in IGN and soma as FPKM. C) Metagene plots of the individual histone marks around target genes of the CSR-1 and ALG-3/4 groups for IGN, oocyte, sperm, early embryo and soma datasets. Normalized signals (as IP/input ratio) were used for plotting. Upstream region (TSS-1000bp), gene body, and downstream region (TES+1000 bp) are included.

To further explore the specific histone marks associated with different sets of AGO targets, and to follow their dynamics during gametogenesis and the onset of embryogenesis, we compared datasets from IGN, oocytes, sperm, early embryos, and soma (Methods). For each timepoint, at least three histone marks were assayed (Figure 4C). Target genes of the CSR-1 group have similarly high levels of H3K4me3 and H3K36me3 in IGN, but H3K36me3 levels specifically drop in gametes and remain low in embryos and soma, while H3K4me3 remains high on these genes. This observation suggests that the CSR-1 small RNA pathway plays a role in reinforcing the H3K36me3 pattern at these genes specifically in the germ line. By comparison, targets of the ALG-3/4 group have low levels of active marks in IGN, but a strong enrichment of H3K4me3 specifically in sperm, suggesting that ALG-3/4 promotes the deposition of this mark during spermatogenesis [48]. These target genes also have elevated H3K27me3 in the soma, especially the targets of RDE-1 and WAGO-10 (Figure 4C), suggesting they are specifically silenced during or after embryogenesis. By contrast, target genes of the ERGO-1 and WAGO-1 groups tend to have relatively low levels of all marks in IGN and gametes, with the exception of H3K9me3, which is particularly high for PPW-1 (Figure S6C). H3K9me3 is detectable in early embryos for targets of all AGOs in the WAGO-1 subgroup, suggesting it persists through gametogenesis, although H3K9me3 profiles are not available for oocytes or sperm. However, in soma, H3K27me3 becomes the dominant silencing mark for these targets, suggesting a major shift during development in the mode of gene silencing at targets of the WAGO-1 group. In sum, gene targets of different small RNA pathways exhibit distinct and dynamic epigenetic profiles in the germline and during development, indicating that post-transcriptional regulation of mRNAs by these pathways is reflected at the level of chromatin.

### H3K4me3 is dynamically redistributed during oogenesis

The dynamic chromatin profiles in IGN, gametes, and embryos at the different target genes of small RNA pathways prompted us to further investigate chromatin remodeling genome-wide across this developmental transition. We first compared the genome-wide distribution of histone modifications in IGN to a published embryo dataset obtained from mixed embryos (Methods). Individual histone modifications were correlated between IGN and mixed-stage embryos (r > 0.20), with the striking exception of H3K4me3 (r = 0.07) (Figure 5A). We first asked whether the low correlation occurred because different genes had H3K4me3 enrichment in the two datasets. IGN have many more genes with significant levels of H3K4me3 than do embryos, indicating extensive remodeling of this state during the germline-to-embryo transition (Figure 5B). Conversely, more than half of the genes associated with H3K4me3 in the embryo were also associated with this mark in IGN, suggesting that maternal H3K4me3 contributes significantly to the embryonic H3K4me3 pattern. Interestingly, the shared genes have a higher transcript abundance than those marked only in IGN or embryo (Figure 4C). Gene Ontology analysis demonstrated that genes marked by H3K4me3 only in IGN or embryos reflect germline or embryogenesis functions respectively, while shared genes represent housekeeping and metabolic genes (Figure S7A).

**Figure 5.**
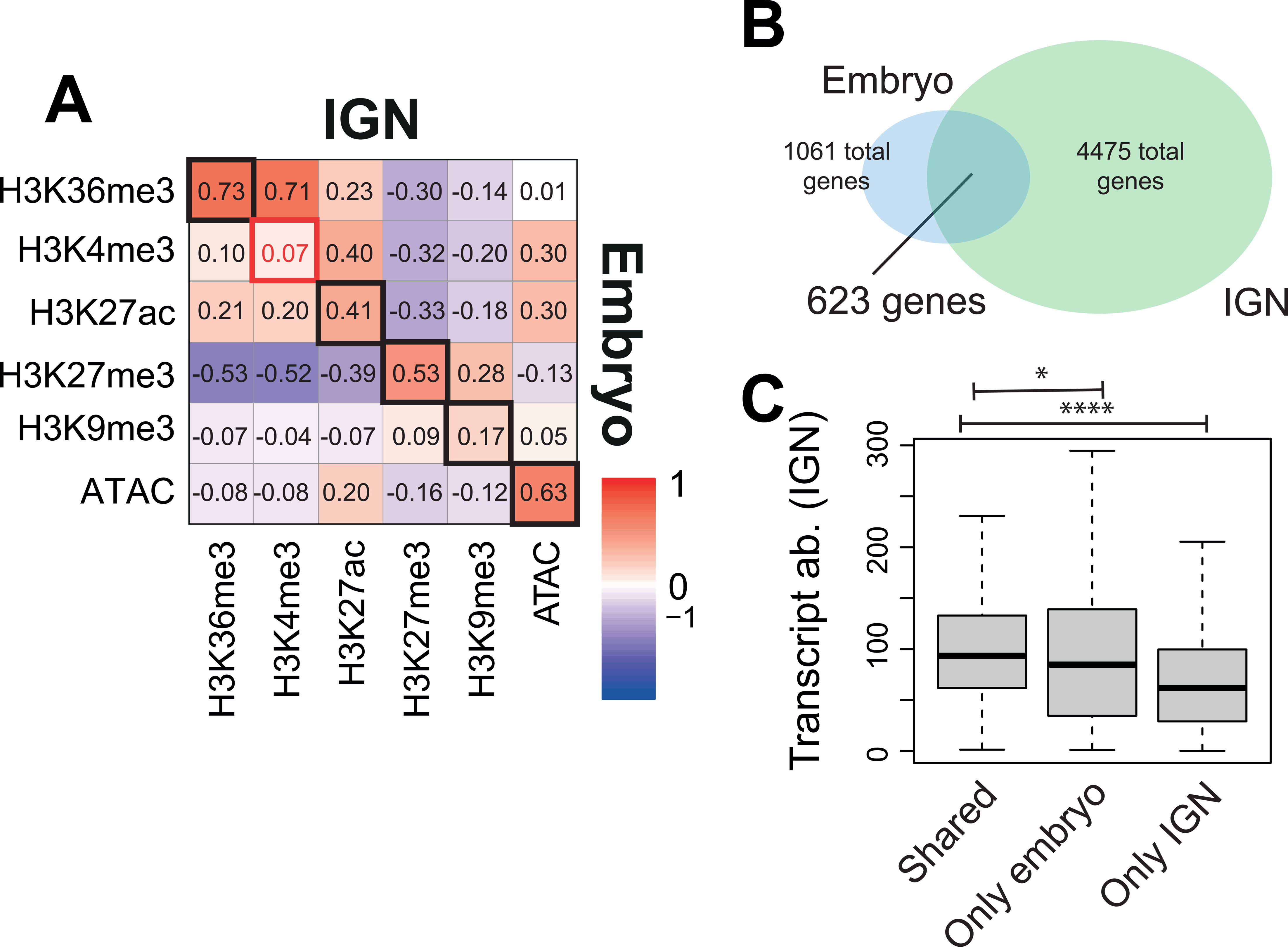
H3K4me3 distribution differs between IGN and embryo. A) Pearson correlation plot of chromatin accessibility for IGN (horizontal) vs. embryo (vertical) for each histone modification. B) Overlap of genes marked by H3K4me3 in IGN vs embryos. Significance assessed by hypergeometric distribution. C) Transcript abundance in IGN for IGN-specific, embryo-specific, and shared genes. P-value legend: * 0.05, ** 0.005, *** 0.0005, **** <0.0005.

H3K4me3 is most often characterized as a relatively narrow peak located at promoters, but has recently been found in a broad pattern during aging in mouse oocytes and also in the soma of *C. elegans* [12,41]. We therefore measured H3K4me3 peak breadth and found that H3K4me3 peaks were significantly broader in IGN than in embryos (Figure 6A). Broad peaks (>3000 bp) were only observed in IGN and represent 35% of total IGN H3K4me3 peaks. Conversely, sharp peaks (<1000 bp) represent 77% of embryonic peaks, but only 27% of IGN peaks. In IGN, these broad H3K4me3 peaks are centered on the gene body, while in the embryo the sharp peaks are located on promoters (Figure 6B). This observation suggests that remodeling H3K4me3 is a global event, independent of individual gene regulation at either developmental stage. We refer to these two patterns as “gene body” and “promoter”, respectively. To more precisely define when these changes occur during the transition between IGN and embryo, we also analyzed H3K4me3 data from isolated oocytes and early embryos [35,5l], which represent intervening developmental timepoints (Figure S7B-D). We observed that the gene body distribution in IGN was gradually restricted to the promoter of a subset of the same genes during this developmental process (Figure 6C). Surprisingly, the shift from the gene body to promoter begins prior to or during oocyte maturation and is subsequently refined during the oocyte-to-embryo transition. To identify which genes underwent this remodeling, we defined sets of genes with high H3K4me3 signal specific to IGN, specific to oocytes, or shared (Figure S6E). As expected, the shared and oocyte-specific genes showed remodeling of H3K4me3, while the IGN-specific genes did not (Figure S6F). GO analysis indicates that the shared and oocyte-specific genes are enriched for housekeeping transcription and translation regulatory factors (Figure S6G). To further probe this observation, we directly asked whether genes with enriched expression in specific tissues were preferentially remodeled in oocytes and early embryos (Figure 6D). H3K4me3 could only be detected for genes with germline or ubiquitous expression, but not for genes specifically expressed in somatic tissues. Together, these analyses indicate that H3K4me3 is remodeled from a gene body pattern in IGN to a promoter-restricted pattern in oocytes at a subset of germline-expressed genes that likely play a role in initiating gene expression in the early embryo.

**Figure 6.**
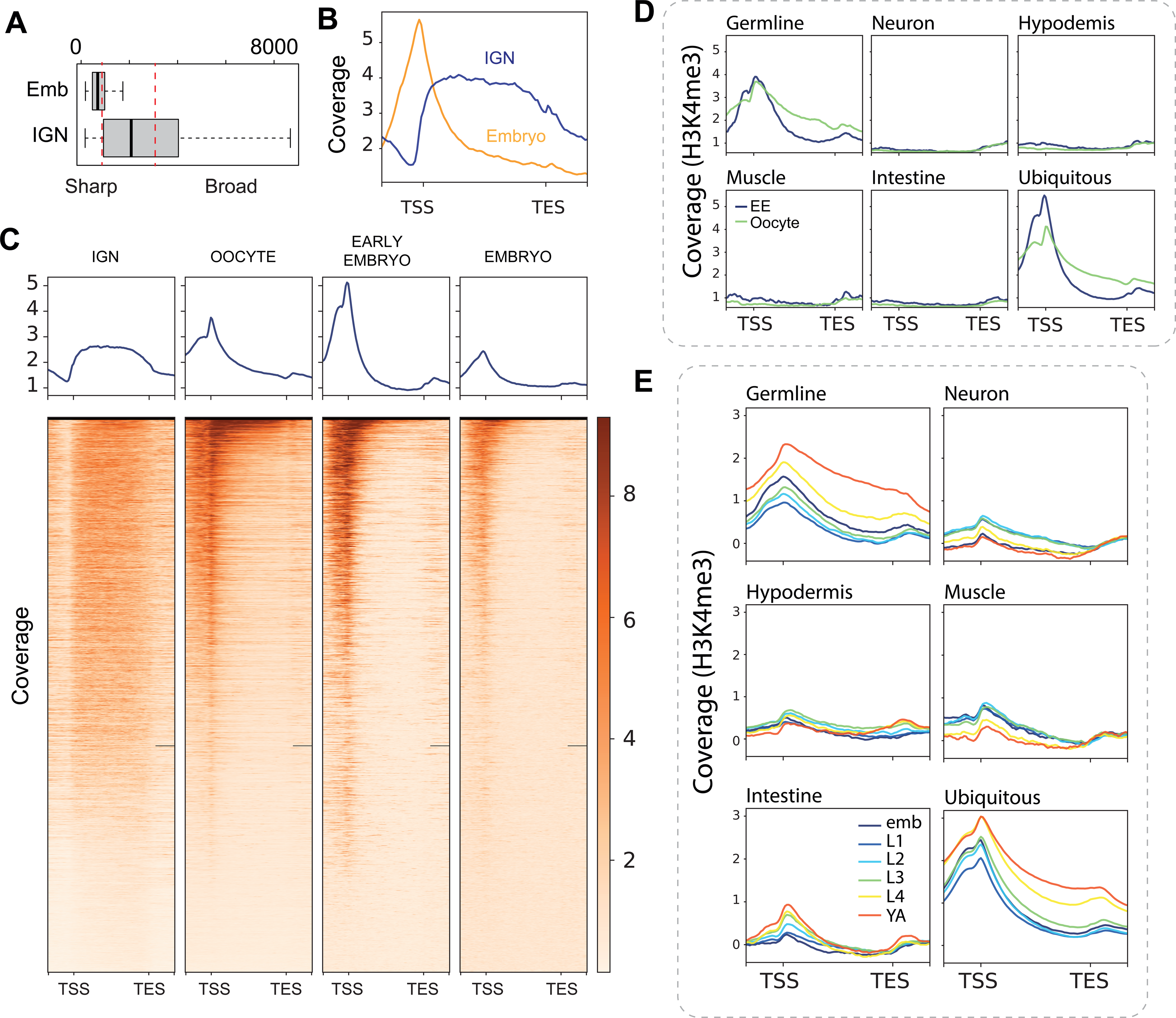
H3K4me3 exhibits dynamic remodeling during oogenesis. A) Breadth of H3K4me3 peaks in embryos (left) and IGN (right). B) Metagene plots of H3K4me3 signal (as normalized IP/input ratio) around genes marked by H3K4me3 in IGN and embryo. C) Metagene plots and heatmaps of H3K4me3 signal in IGN [23], oocytes [51], early embryos [51], and mixed embryos [25]. D) Metagene plots of H3K4me3 signal in early embryos and oocytes for genes with tissue-specific expression [47]. E) Metagene plots of H3K4me3 signal during larval development for genes with tissue-specific expression. Normalized IP/input ratio is plotted. Upstream region (TSS-1000bp), gene body, and downstream region (TES+1000 bp) are included.

In *C. elegans*, the broad H3K4me3 domain has been related to the aging process in somatic tissues, suggesting it might be present in adult IGN as an indirect consequence of development or aging [41]. We therefore asked whether the broad H3K4me3 domain occurred specifically in IGN or was found in somatic cells as well, by analyzing H3K4me3 profiles at each developmental stage from whole worms [25], segregated by sets of genes with tissue-specific expression [47]. Strikingly, the H3K4me3 pattern shifted from promoter to gene body over development for germline-specific and ubiquitously-expressed genes, which are also expressed in the germline, but not for genes expressed specifically in somatic tissues (Figure 6E). These observations indicate that the appearance of broad H3K4me3 peaks on gene bodies likely occurs primarily in the germ line. To confirm that this remodeling relative to gene structure is specific to H3K4me3, we compared its profile to H3K27ac, which is also associated active gene expression and usually restricted to promoters. Unlike H3K4me3, H3K27ac has a promoter-restricted distribution on approximately the same percentage of genes in each stage (Figure S7H), indicating this remodeling is indeed unique to H3K4me3. In sum, we have characterized a major shift in H3K4me3 distribution that occurs during oocyte maturation, before the oocyte-to-embryo transition on genes that are expressed in the germ line.

## DISCUSSION

In this report, we have collected and analyzed epigenetic data from diverse genomic assays to characterize the overall chromatin state of adult hermaphrodite germ cells in *C. elegans*. The distribution of chromatin states across the genome captures known and novel features of genome organization in the germ line, and identifies germline-specific chromatin states. Distinct chromatin states at various sets of germline-expressed genes point to feedback from post-transcriptional regulatory mechanisms, including small RNA pathways, into epigenetic state. In addition, we found that chromatin state dynamics during the oogenesis-to-embryogenesis transition revealed that most histone modifications remain fairly stable, with the exception of H3K4me3. We have made these chromatin states available for further computational analysis and for viewing via genome browser (File S1). This analysis sets the stage for further investigation into the underlying mechanisms that establish these states, and the phenotypic consequences to germline development and embryogenesis when these states are disrupted.

Several unique features of germline gene regulation were apparent in this analysis, and certain differences in chromatin states between the germ line and soma suggest distinct regulatory mechanisms. For instance, the “strongly inactive TSS” state 5 found only in IGN is characterized by conflicting signals of open chromatin that is marked at the promoter by the silencing modification H3K27me3 (Figure 1, 2). To a much lesser extent, state 7 “primed TSS” also has a similar profile. Genes associated with these states tend to function in signal transduction, neuronal activity, and development (Figure S1C), and mostly do not show enriched expression in the germ line. Thus, they might functionally approximate the “poised” chromatin state found in germ cells of other organisms, in which genes expressed in the embryo are placed in a chromatin state in the germline that prevents immediate transcription but facilitates rapid expression post-fertilization [36]. However, the chromatin hallmarks of poising in mammals are the co-localization of the opposing marks H3K27me3 and H3K4me3 [36], a configuration not found in any germline chromatin state in *C. elegans*. Thus, this observation suggests the existence of an alternate mechanism that employs only H3K27me3 to temporally delay gene expression across the oocyte-to-embryo transition.

Additionally, certain chromatin states showed complex correlations between sites of open chromatin, accumulation of multiple active histone marks, and transcript abundance. A priori, one would expect that transcript abundance would be roughly proportional to the level of active histone marks and chromatin accessibility, as seen overall for the germline chromatin states (Figure 1A, B). However, we found multiple chromatin states with higher or lower transcript abundance than might be expected based on the combined levels of active/inactive modifications (e.g. state 1 - pregamete, state 9 – spermatogenesis, state 11 – oogenesis, Figure 3). These instances suggest the existence of extensive post-transcriptional regulation that stabilizes or degrades transcripts independently of chromatin state. Indeed, many mechanisms of post-transcriptional regulation in the germ line have been demonstrated to affect the mitosis-to-meiosis transition, the sperm-to-oocyte switch, and other critical aspects of germline development [18,49].

In particular, post-transcriptional regulation through small RNA pathways not only affects target mRNA transcripts, but can lead to alterations of chromatin modifications at the genomic locus of those targets. The best-studied example is the AGO protein HRDE-1, which localizes to the nucleus in germ cells and promotes deposition of H3K9me3 at target loci [48]. Other AGO proteins might also have a similar effect, either directly or indirectly, although this possibility has not been thoroughly investigated. Indeed, we do see enrichment of H3K9me3 at HRDE-1 target genes, but this enrichment is actually much stronger for the targets of PPW-2, which is in the same WAGO-1 group as HRDE-1 but does not exhibit detectable nuclear localization (Figure S7C). Additionally, AGOs in the ALG-3/4 group, which promote spermatogenesis [48], display a striking increase of H3K4me3 at target genes specifically in sperm, suggesting a downstream effect of ALG-3/4 on the chromatin state of its targets. Finally, targets of the CSR-1 group exhibit high levels of the active marks H3K36me3 and H3K4me3 in IGN, as expected since these AGOs target germline-expressed genes. However, for all these genes, H3K36me3 is primarily present only in the germ line and begins to decrease in gametes and in embryos, while H3K4me3 persists throughout the oocyte-to-embryo transition and into later somatic development (Figure 6C). This observation raises the possibility that CSR-1 is functionally linked to H3K36me3 at its targets in the germline. Intriguingly, CSR-1, which is perinuclear throughout most of germline differentiation, enters the nucleus in oocytes and early embryos [10], when the H3K36me3 levels are declining. CSR-1 might therefore play a direct role in altering this modification during this transition. The mechanisms by which most AGO proteins affect these marks directly or indirectly is currently unclear and will require functional investigation of many of the different histone methyltransferases and demethylases expressed in the germ line.

One of the most striking observations of this analysis is the extensive remodeling of H3K4me3 during the oocyte-to-embryo transition (Figure 4, 5). This remodeling is two-fold: the genes enriched for with H3K4me3 change, and the specific distribution pattern of H3K4me3 across genes changes from a broad “gene body” pattern to a focused accumulation at promoters. H3K4me3 has been primarily described as a promoter mark, but recent studies have found instances in which it covers larger genomic regions including during aging in *C. elegans* [12,41]. H3K4me3 is also remodeled from broad peaks to promoter peaks during the mouse maternal-to-zygotic transition [12], but the functional consequence of such remodeling is unclear, as is the underlying mechanism. Intriguingly, a recent report demonstrated that the canonical histone H3 is replaced with the variant H3.3 at the approximate time of this H3K4me3 remodeling, during oogenesis [21]. H3.3 is the dominant form specifically during early embryogenesis, until it is again replaced by canonical H3 later in development. This time frame corresponds with the H3K4me3 remodeling that we report here, suggesting a possible connection. Identifying the regulators of H3K4me3 distribution remodeling during the oogenesis-to-embryogenesis transition will be critical next steps in understanding its functional consequences.

This integrated analysis of chromatin state in the *C. elegans* germ line establishes an initial overview that provides insight into gene regulatory processes and germline biology. However, this approach is limited in some aspects. The catalog of histone modifications is far from complete, and the included genomic assays are restricted to linear chromatin so that any correlations between the chromatin state and three-dimensional organization in the germline are unknown. Additionally, while the use of IGN greatly improves specificity and resolution, it represents an aggregate picture of millions of nuclei at many different points along the germ cell differentiation trajectory. In the future, technologies that simultaneously measure chromatin accessibility and RNA levels in single cells would help to further clarify the dynamics of chromatin state during germ cell differentiation. Regardless, characterizing adult IGN chromatin states using multiple histone modifications and chromatin accessibility, as we have done here, provides a powerful foundation on which to pursue finer-detailed characterizations and more mechanistic studies.

## ACKNOWLEDGEMENTS

This work was funded by NIH R35GM131776.

---

**Supplemental Figure 1.**
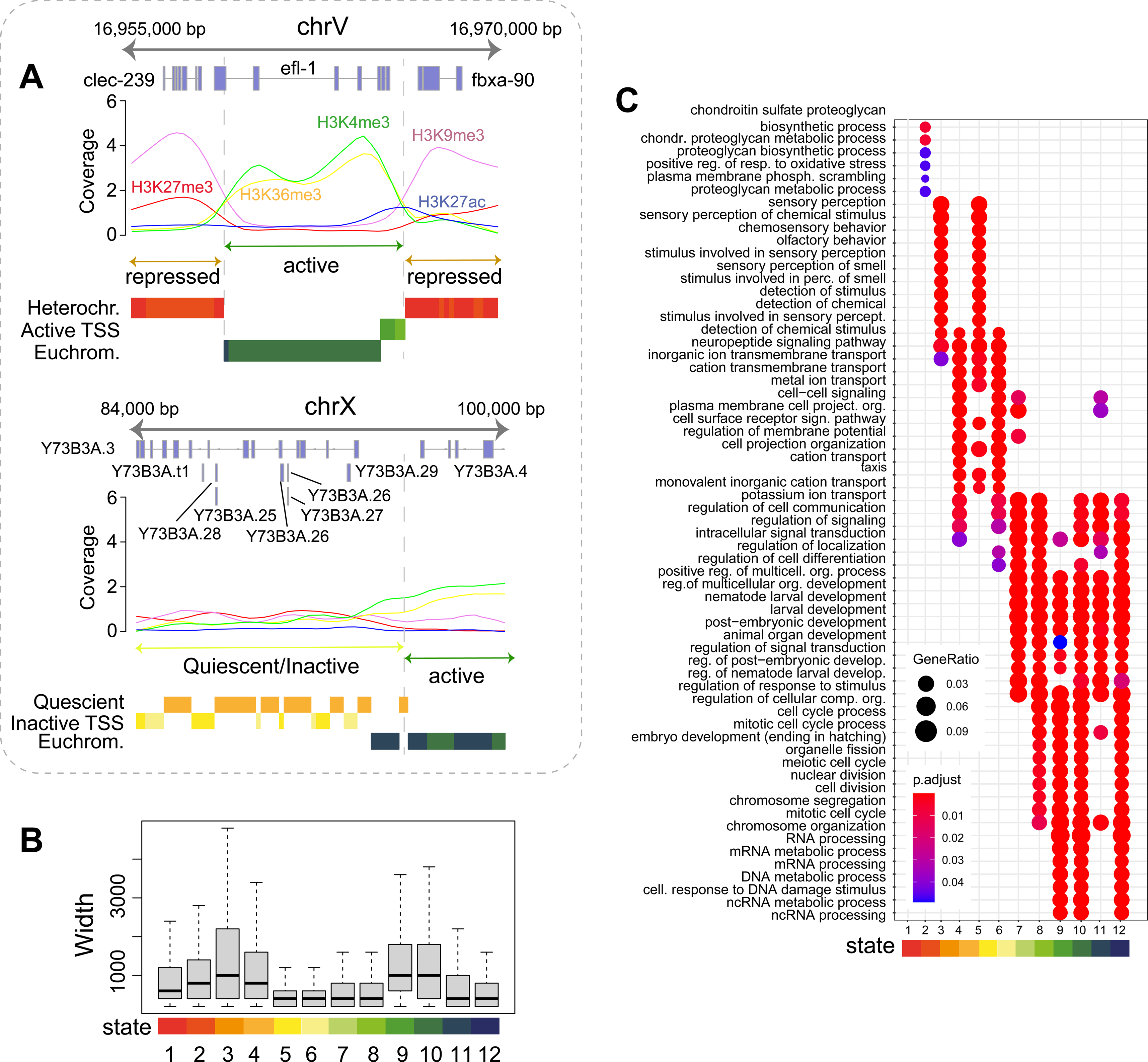
Characteristics of germline chromatin state. A) Genomic visualization of the chromatin states along a portion of chromosome V (above) and chromosome X (below). B) Width in base pairs of the genomic sections for each state. C) Enriched categories for genes contained in each chromatin state using Gene Ontology analysis; only significantly enriched categories (with adjusted p-value < 0.1) were plotted as a dotplot. Both gene ratio and adjusted p-values are included as shape size, and color code respectively.

**Supplemental Figure 2.**
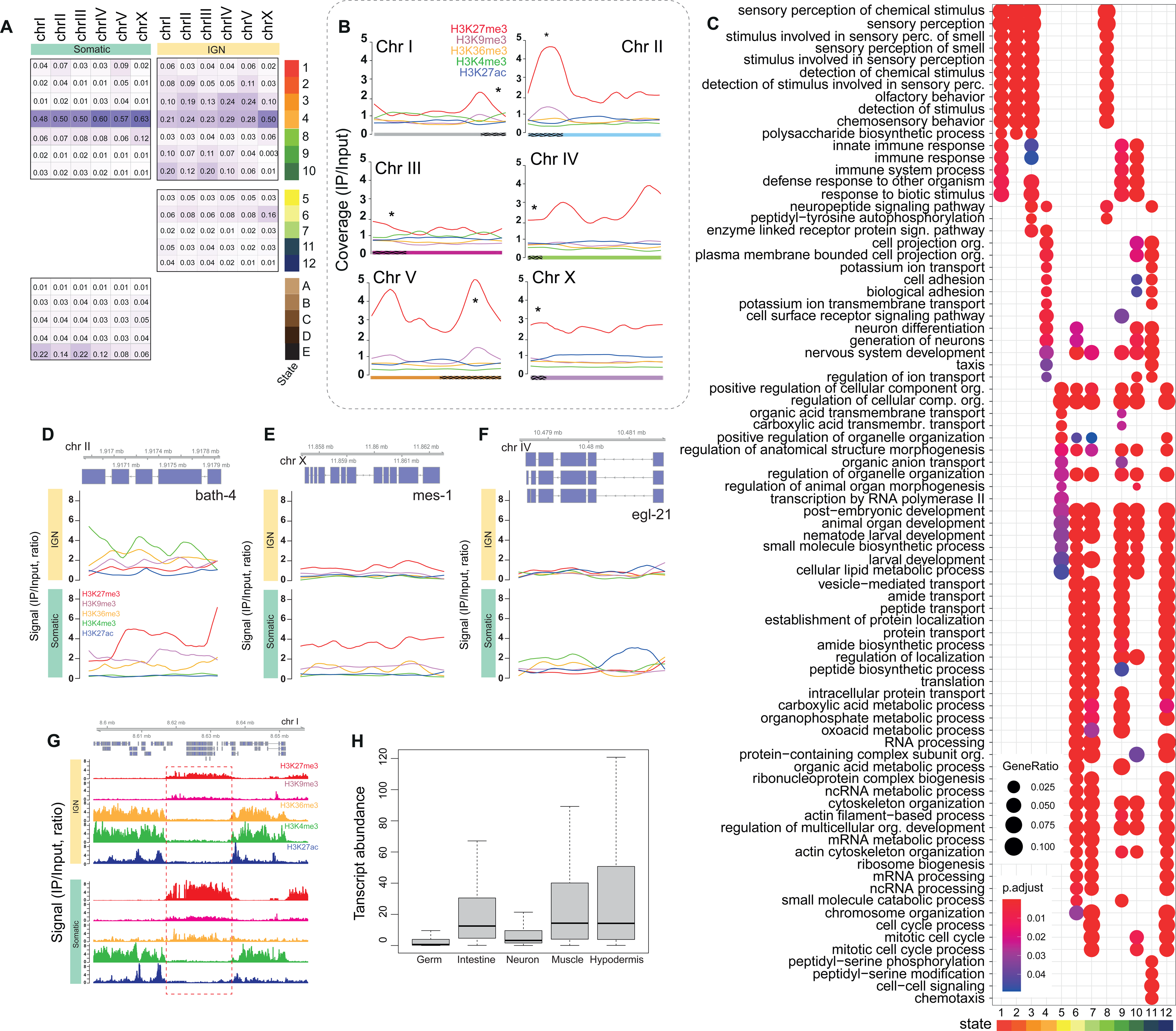
Chromatin states unique to germ line and soma. A) Percentage of the individual chromosomes in each chromatin states for germline and somatic tissues. Chromatin states are divided in “shared” states (above), “only-IGN” states (center), and “only-som” states (below). B) Distribution of normalized signal (as IP/input ratio) for each epigenetic mark along each chromosome in the soma. C) Enriched GO categories for genes contained in each somatic chromatin state; only significantly enriched categories (with adjusted p-value < 0.1) were plotted as a dotplot. Both gene ratio and adjusted p-values are included as shape size, and color code respectively. D-F) Examples of genes that go through an “epigenetic” switch from germline to somatic data: *bath-4* (D), *mes-1* (E), and *egl-21* (F). The genomic portion of the chromosome, and the annotated genes, are visualized. G) Genomic visualization of the chromatin states along a portion of chromosome I shows additional epigenetic differences between germ line and soma. The genomic portion of the chromosome, and the annotated genes, are visualized. H) Transcript abundance of tissue-specific genes [46] in somatic data.

**Supplemental Figure 3.**
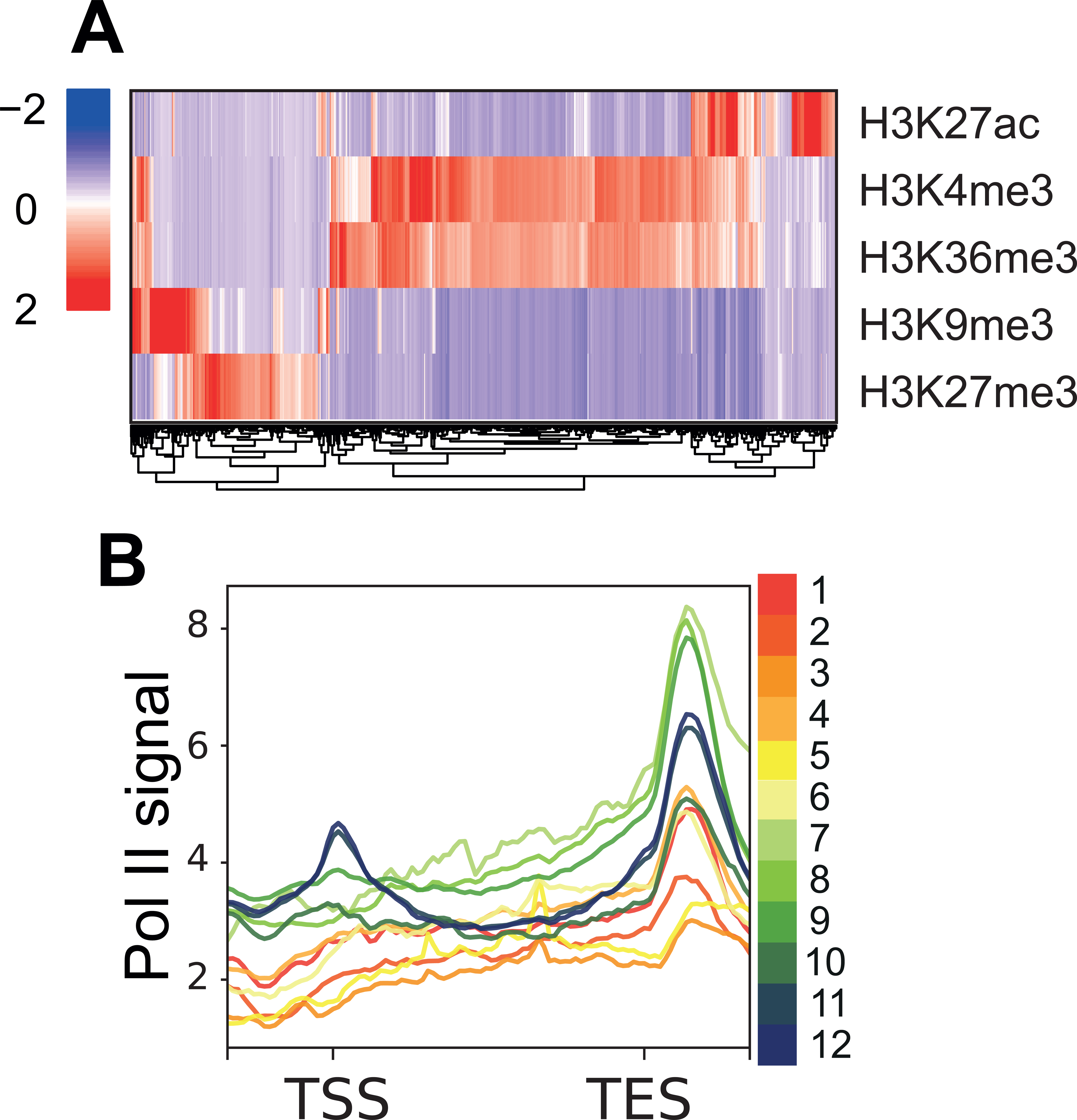
Chromatin states of classes of germline-expressed genes in IGN - pregamete. A) Heatmap of the individual histone modification for genes with pregamete-enriched expression. Values are centered on the mean and scaled on the standard deviation. B) Polymerase II signal (whole worms) for genes with pregamete-enriched expression in each state. Upstream region (TSS-1000bp), gene body, and downstream region (TES+1000 bp) are included.

**Supplemental Figure 4.**
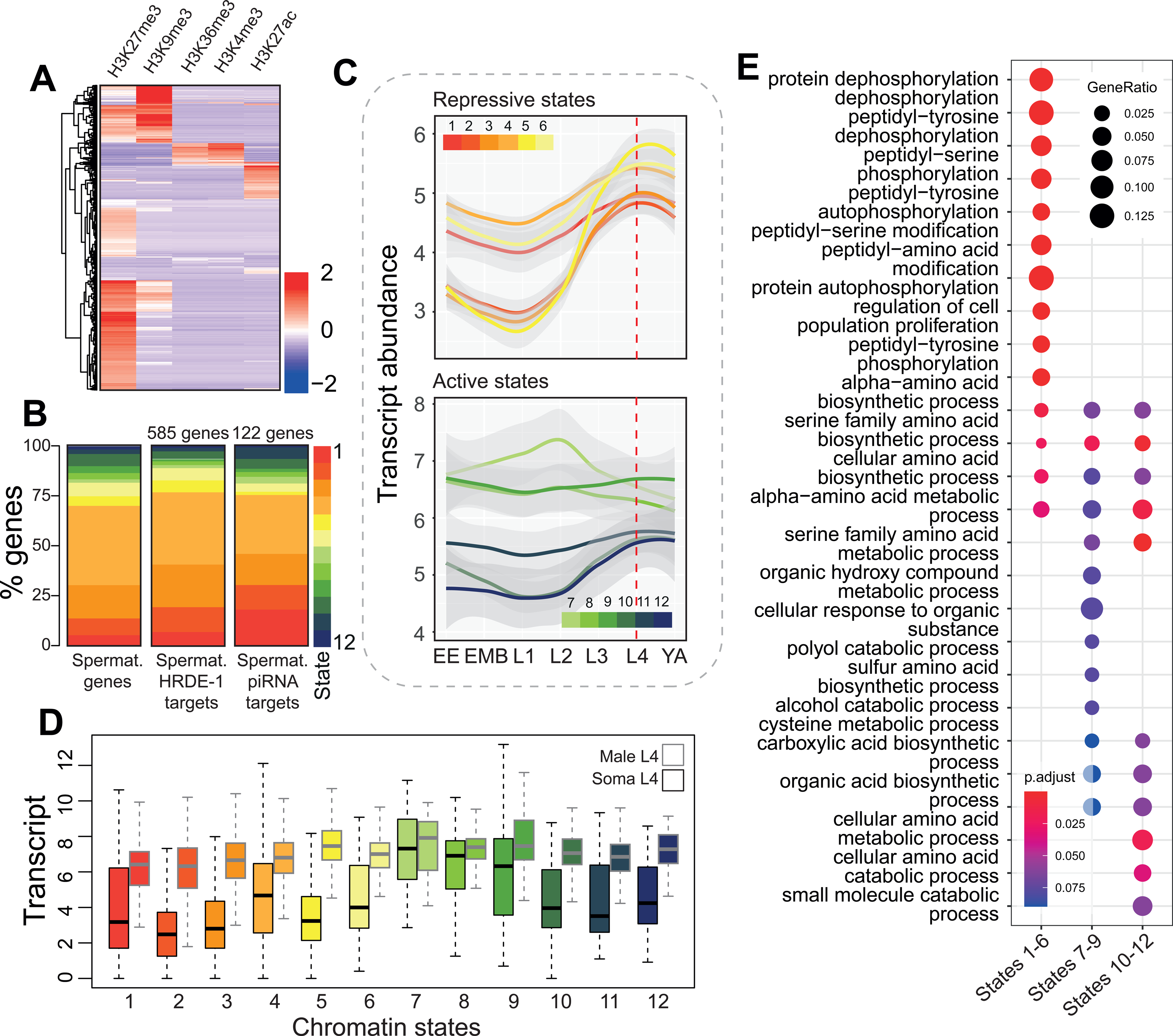
Chromatin states of classes of germline-expressed genes in IGN - spermatogenesis. A) Heatmap of the individual histone modification for genes with spermatogenesis-enriched expression. Values are centered on the mean and scaled on the standard deviation. B) Distribution of chromatin states for all genes with spermatogenesis-enriched expression (left), only those that are targets of HRDE-1 (middle), and only those that are piRNA targets (right) [11]. C) Gene expression developmental profiles for genes with spermatogenesis-enriched expression in each state using transcriptomic data [7]. Log normalized counts are visualized. D) Gene expression profiles for genes with spermatogenesis-enriched expression in each state in somatic tissues (black outline) and male (grey outline). E) Gene Ontology analysis of genes with spermatogenesis-enriched expression for combined chromatin states - inactive (states 1-6), expressed (states 7-9), and active but not expressed (states 10-12). Both adjusted p-value and gene ratio are included.

**Supplemental Figure 5.**
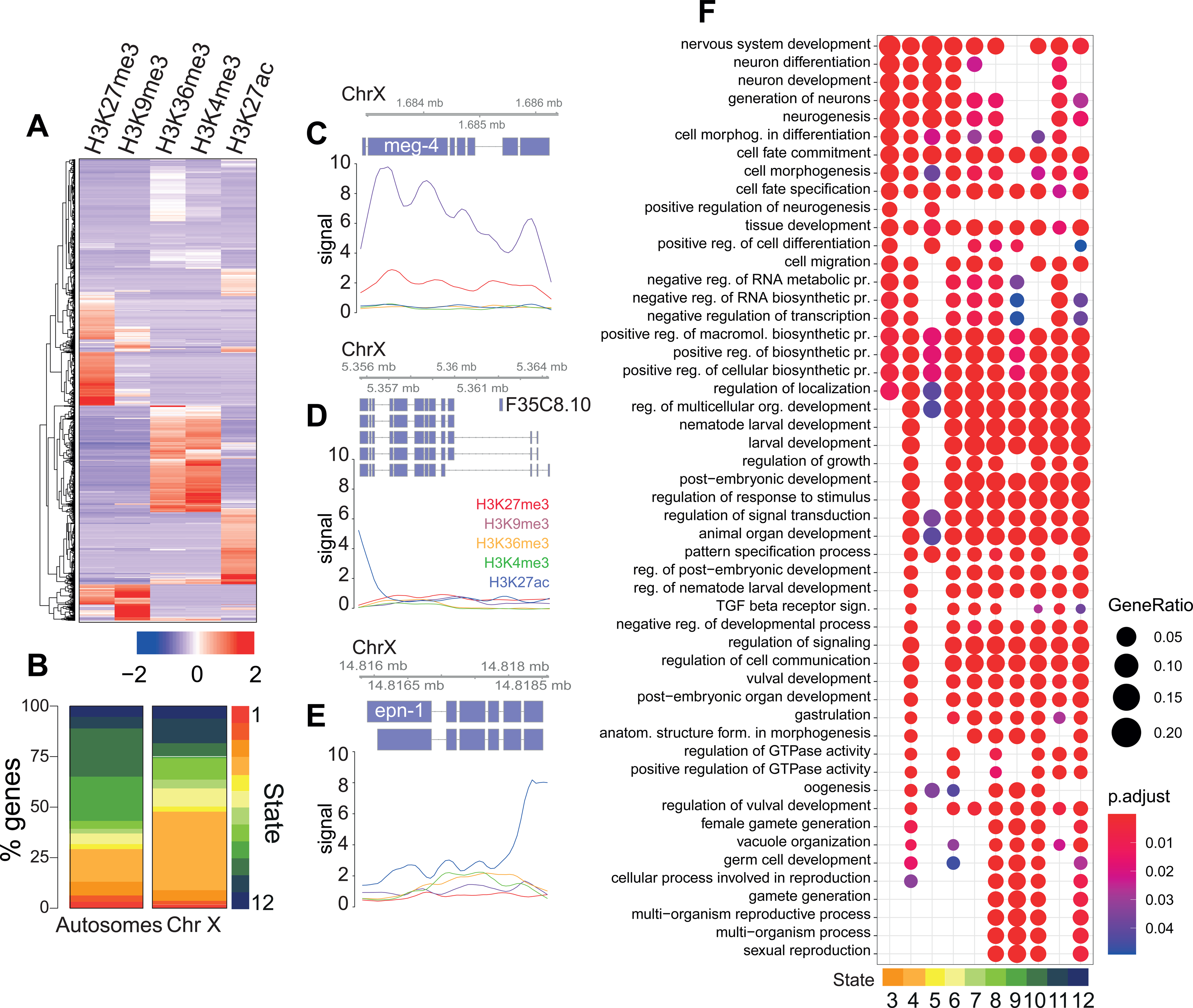
Chromatin states of classes of germline-expressed genes in IGN – oogenesis. A) Heatmap of the individual histone modification for genes with oogenesis-enriched expression. Values are centered on the mean and scaled on the standard deviation. B) Chromatin states for oogenesis-enriched genes on autosomes and chromosome X. C-E) Genomic visualization of histone mark levels along *meg-4* (C), *chtl-1* (D), and *epn-1* (E). The genomic portion of the chromosome, the annotated genes, as well as the related chromatin state, are visualized. F) Gene Ontology analysis of genes with oogenesis-enriched expression for chromatin states 3-12. Adjusted p-value and gene ratio are included.

**Supplemental Figure 6.**
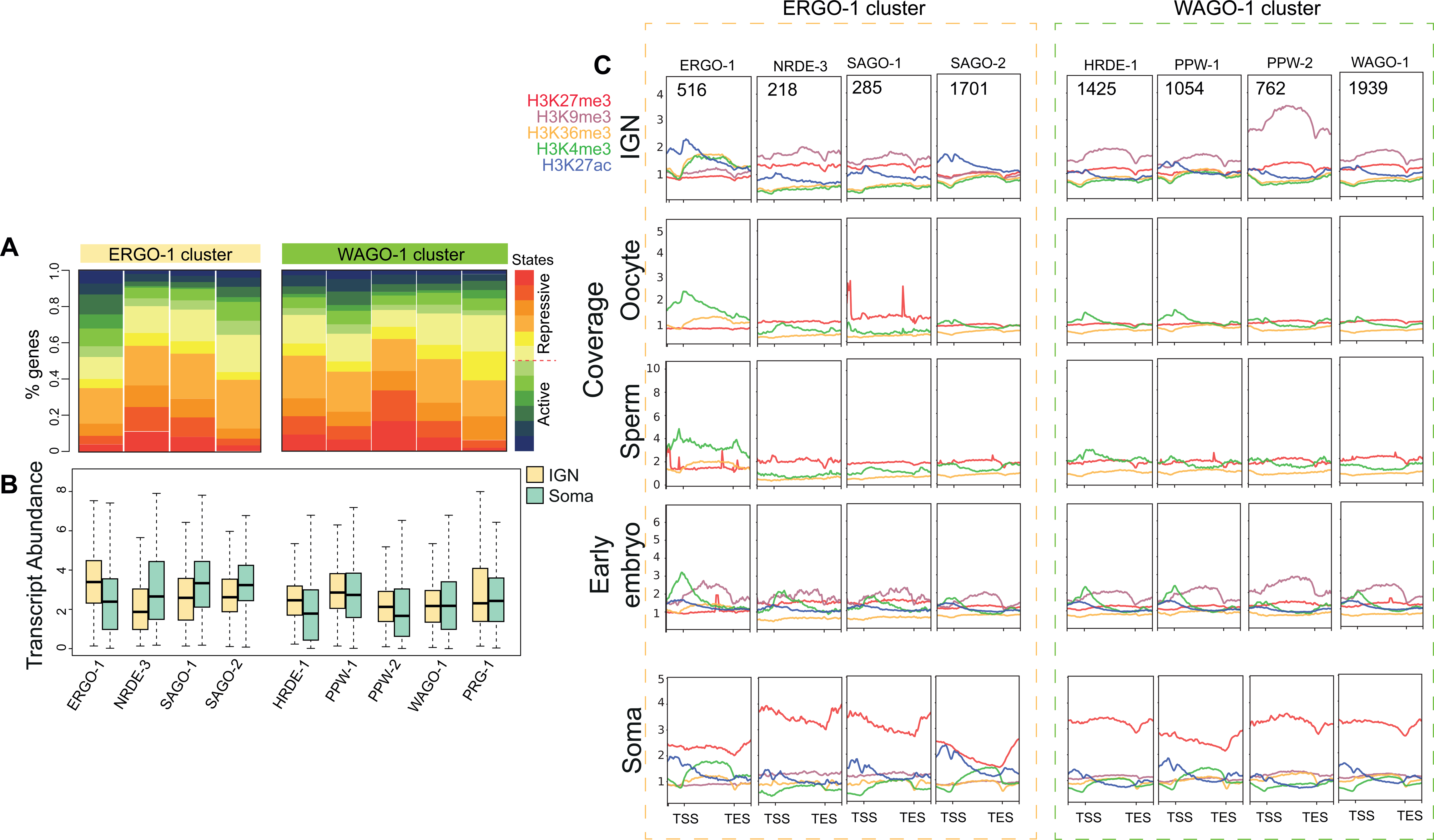
Chromatin states of AGO target genes. A) Distribution of chromatin states along targets of AGO proteins in the ERGO-1 and WAGO-1 groups [48]. B) Transcript abundance (FPKM) of these targets in IGN and soma. C) Metagene plots of the individual histone marks around target genes of the ERGO-1 and WAGO-1 groups for IGN, oocyte, sperm, early embryo and soma datasets. Normalized signals (as IP/input ratio) were used for plotting. Upstream region (TSS-1000bp), gene body, and downstream region (TES+1000 bp) are included.

**Supplemental Figure 7.**
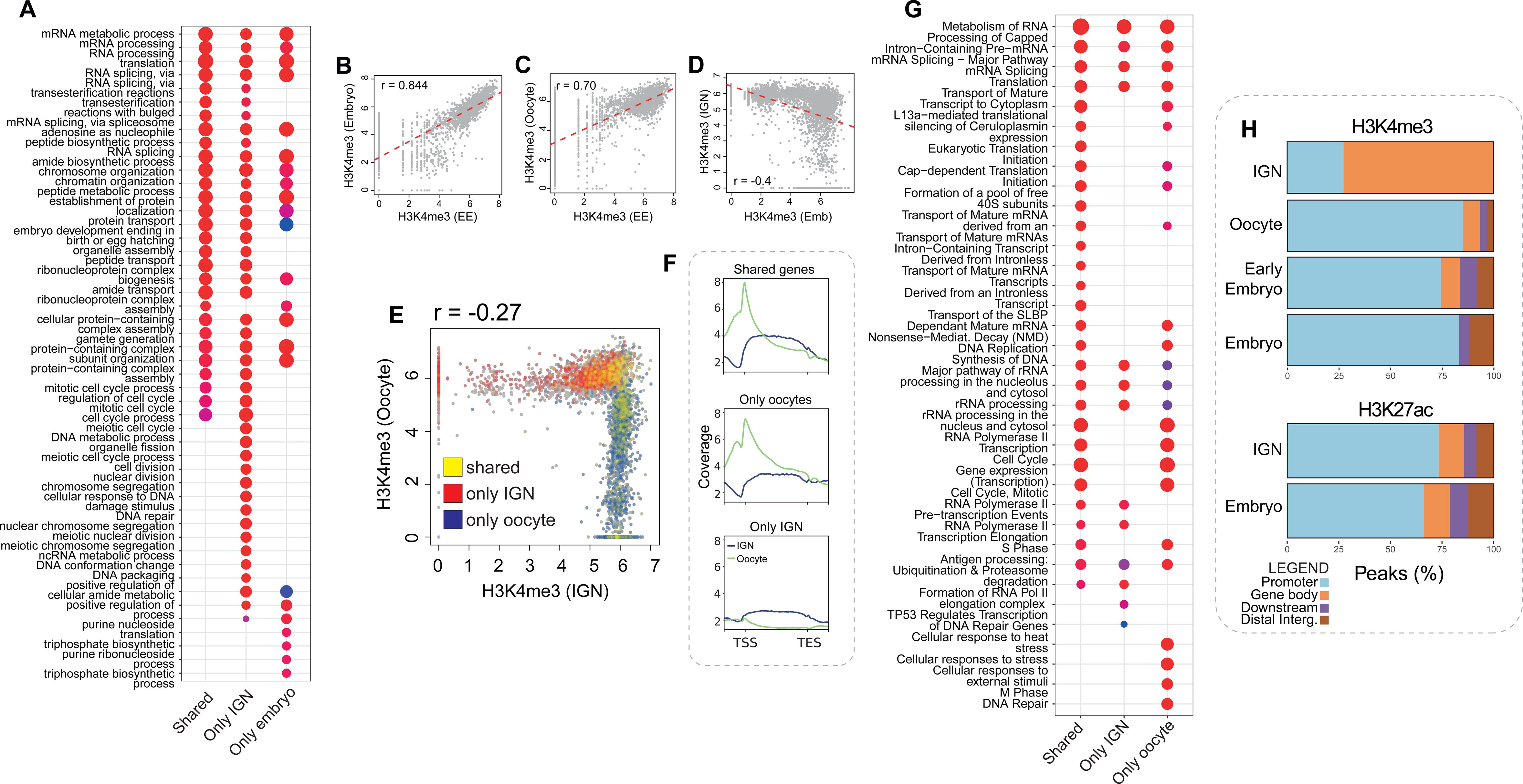
H3K4me3 exhibits dynamic remodeling during oogenesis. A) GO analysis of genes marked by H3K4me3 in both IGN and embryos (“shared”), genes marked only in IGN (“Only IGN”), and genes marked only in embryos (“Only embryo”). Adjusted p-value and gene ratio are included. B-D) Correlation plots of H3K4me3 signal (as normalized IP/input log ratio) between different stages of development, in 500bp bins. Pearson correlation and slope are also shown. B. Embryo (X-axis) vs oocyte (Y-axis), C Embryo vs early embryo, D. Embryo vs IGN. Embryo data from [25], oocyte and early embryo from [51], and IGN from [23]. E) Plot of H3K4me3 signal between IGN and oocyte as in B-D. Genes marked by H3K4me3 in both IGN and oocytes (yellow), genes marked by H3K4me3 only in oocytes (red), and genes marked by H3K4me3 only in IGN (blue) are highlighted. F) Metagene plots of H3K4me3 signal (as normalized IP/input ratio) around genes marked by H3K4me3 in both IGN and oocytes, only in IGN, and only in oocytes. Upstream region (TSS-1000bp), gene body, and downstream region (TES+1000 bp) are included. G) GO analysis of genes marked by H3K4me3 in both IGN and oocytes (“shared”), genes marked only in IGN (“Only IGN”), and genes marked only in oocytes (“Only oocyte”). Both adjusted p-value and gene ratio are included. H) Distribution of H3K4me3 and H3K27ac across gene features at multiple developmental stages.

## Notes

### Competing Interest Statement

The authors have declared no competing interest.

